# A Clustering Approach for Detecting Implausible Observation Values in Electronic Health Records Data

**DOI:** 10.1101/570564

**Authors:** Hossein Estiri, Shawn N. Murphy

## Abstract

**Background:** Identifying implausible clinical observations (e.g., laboratory test and vital sign values) in Electronic Health Record (EHR) data using rule-based procedures is challenging. Anomaly/outlier detection methods can be applied as an alternative algorithmic approach to flagging such implausible values in EHRs.

**Objective:** The primary objectives of this research were to develop and test an unsupervised clustering-based anomaly/outlier detection approach for detecting implausible observations in EHR data as an alternative algorithmic solution to the existing procedures.

**Methods:** Our approach is built upon two underlying hypotheses that, (i) when there are large number of observations, implausible records should be sparse, and therefore (ii) if these data are clustered properly, clusters with sparse populations should represent implausible observations. To test these hypotheses, we applied an unsupervised clustering algorithm to EHR observation data on 50 laboratory tests. We tested different specifications of the clustering approach and computed confusion matrix indices against a set of silver-standard plausibility thresholds. We compared the results from the proposed approach with conventional anomaly detection (CAD) approach’s, including standard deviation and Mahalanobis distance.

**Results:** We found that the clustering approach produced results with exceptional specificity and high sensitivity. Compared with the conventional anomaly detection approaches, our proposed clustering approach resulted in significantly smaller number of false positive cases.

**Conclusion:** Our contributions include (i) a clustering approach for identifying implausible EHR observations, (ii) evidence that implausible observations are sparse in EHR laboratory test results, (iii) a parallel implementation of the clustering approach on i2b2 star schema, and (3) a set of silver-standard plausibility thresholds for 50 laboratory tests that can be used in other studies for validation. The proposed algorithmic solution can augment human decisions to improve data quality. Therefore, a workflow is needed to complement the algorithm’s job and initiate necessary actions that need to be taken in order to improve the quality of data.

## 1. Introduction

Data stored in Electronic Health Records (EHR) offer promising opportunities to advance healthcare research, delivery, and policy. Provision of these opportunities is contingent upon high quality data for secondary use. Data quality concerns, however, have hampered secondary use of EHR data.[1,2] The increasing throughput of EHR data constantly deposited into clinical data research networks have cultivated new opportunities for utilizing innovative statistical learning methods to improve quality.

Plausibility is a dimension of data quality that represents whether EHR data values are “believable” or “truthful.”[3] It is quite possible to witness an implausible observation (IO) in EHR data, such as a negative A1c value. An IO is extremely unlikely to signify a fact about a patient and may represent an underlying data quality issue. Detecting such IOs in EHR data are difficult for two reasons. First, gold standards are not always available for all clinical observations to set cut-off thresholds for an implausible observation. Second, even if gold standards are present, detection of out-of-range observations would require manual (rule-based) procedures, which can become increasingly extensive given the diversity of observations, ontologies and measurement units across institutions.

With the abundance of unlabeled data (e.g., vital signs) in EHR repositories, unsupervised learning can offer solutions for characterizing clinical observations into meaningful sub-groups. In unsupervised learning, the machine develops a formal framework to build representations of the input data to facilitate further prediction and/or decision making.[4] We focus on laboratory result records in the EHR. Multiplicity of laboratory tests in EHRs makes it extremely difficult to manually assign rules for identifying implausible values for each lab test. Conceptualizing implausible laboratory test results and vital sign values in EHRs as outliers, we propose and evaluate the feasibility of an unsupervised clustering approach for detecting implausible EHR vital sign and laboratory test values in large scale clinical data warehouses.

## 2. Materials and Methods

Outlier detection a rich area of research in data mining[5,6] and has been extensively applied to clinical data for addressing different issues, such as detecting unusual patient-management actions in ICU,[7] deriving workflow consensus from multiple clinical activity logs,[8] characterizing critical conditions in patients undergoing cardiac surgery,[9] discovering unusual patient management,[10] alert firing within Clinical Decision Support Systems,[11] finding clinical decision support malfunctions,[12] identifying high performers in hypoglycemia safety in diabetic patients,[13] and classifying the influence factor in diabetes symptoms.[14]

There are many algorithms developed to detect outliers based on different approaches to what constitutes an outlier, for which there is no universally agreed definition.[5] Generally, outlier detection refers to the problem of discovering data points in a dataset that do not conform to an expected exemplar.[15]

Outlier detection methods can be characterized into two broad groups of parametric and non-parametric approaches. Parametric methods [16,17] detect outliers by comparing data points to an assumed stochastic distribution model. In contrast, non-parametric (model-free) techniques do not assume a-priori statistical model. Non-parametric methods range in computational and implementation complexity. For example, simple methods such as Grubbs’ method (i.e., Extreme Studentized Deviate) [18] and informal boxplots to visually identify outliers [19] are conveniently used when datasets do not hold complex patterns. Parametric and non-parametric density-based methods for outlier detection are popular among researchers.[20] These methods learn generative (stochastic) density models from data and then identify data points with low probabilities as outliers. For example, Gaussian Mixture Models (GMMs) have been applied to anomaly detection problems. Human biological data have varying density functions that reduce the utility of using parametric density-based methods.

Proximity-based methods are also popular outlier detection techniques that are often simple to implement. Proximity-based methods are categorized into distance- and density-based techniques. In distance-based methods, an outlier is far away from its nearest neighbors (based on local distance measures).[21,22] Mahalanobis distance and *χ*^2^test are popular outlier detection techniques for multivariate data. Distance-based methods can often handle large datasets.[6] However, proximity-based methods suffer from computational intensity in highly dimensional data,[5] and identifying the proper distance measure to identify outliers in challenging. In human biological data (e.g., vital signs and labs data), distance-based methods can lead to high false positive rates.

Clustering-based outlier detection techniques also apply different clustering methods to identify sparse clusters or data points as outliers.[23]

### 2.1. Approach

We propose an unsupervised clustering approach to detect outlier values in lab tests and vital sign values based on two hypotheses. First, we hypothesize (hypothesis #1) that implausible observations – that are not due to systematic human error – must be sparse (infrequent) in datasets that contain large amounts of data. For example, we do not expect to frequently see a blood pressure record of 1090/80 in an EHR repository. Given ontological harmony, we can use data from EHRs to compare an individual data point with a large set of similar data points and identify implausible observations. Therefore, we also hypothesize (hypothesis #2) that a well-specified unsupervised clustering algorithm should be able to partition clinical observations into meaningful clusters, from which we can extract clusters with sparse populations (very small members of data points) as implausible observations.

Figure 1 illustrates the rationale behind our second hypothesis. The bell-shaped curve in Figure 1 is a probability density function for a laboratory result in EHR – extracted and visually modified from Cholesterol in HDL. The hypothetical thresholds for normal ranges and implausible values are delineated on the plot. An unsupervised clustering algorithm partitions the lab values into *n* cluster, each of which embrace a number of data points. We can obtain the number of data points in each cluster and flag clusters with population smaller than a certain threshold as anomalies. Unlike conventional methods such as using standard deviation to identify anomalies, the clustering approach would provide a more flexible solution that is density-based. In addition, the clustering approach should be able to detect implausible observations regardless of their values.

**Figure 1.**
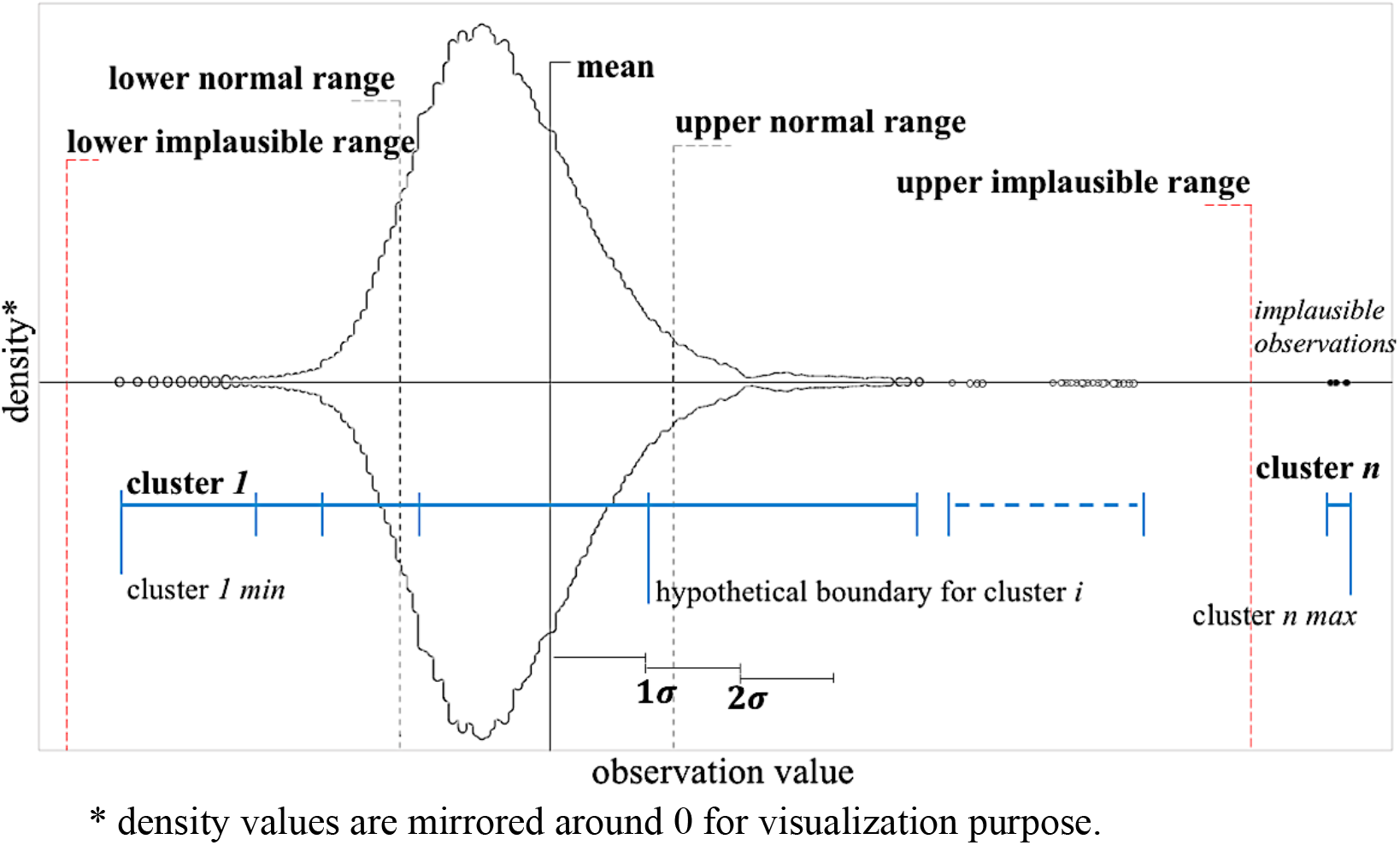
Detecting implausible observations through unsupervised clustering.

To test these hypotheses, we implemented the clustering solution on EHR observation data, using a hybrid hierarchical K-means clustering algorithm. We set different ratios for flagging a cluster as anomalous. To compare the results against conventional methods, we also performed anomaly detection using conventional approaches (CAD) using standard deviation and Mahalanobis distance[25,26] with different configurations for each.

We measured the performance for each algorithm against a set of silver-standard high and low implausible thresholds that we manually curated based on expert judgment, insight from data distributions, and literature search – Table of silver standards is available in the Appendix.

### 2.2. Data

The labs data contained upwards of 720 million rows of data representing 50 laboratory observations with distinct Logical Observation Identifiers Names and Codes (LOINC). On average, each lab observation contained more than 14 million data points, ranging from 45,000 to above 69 million.

### 2.3. The HK-means algorithm

The goal in unsupervised clustering is to partition data points into clusters with small pairwise dissimilarities.[27,28] K-means [29] is one of the most popular unsupervised clustering algorithms,[27] for its simplicity and efficiency. It is a top-down algorithm that aims to minimize the distortion measure by iteratively assigning data points to cluster centroids to meet a convergence criterion.[4,27] K-means is sensitive to outliers.[30,31] Although this property is often considered as a weakness, sensitivity to outliers makes K-means a good algorithm for our purpose of identifying rare events in clinical observations.

For a vector of observations *x*(1), *x*(2),…, *x*(*n*), where *x*(*i*) ∈ ℝ, the K-means algorithm aims to predict *k* centroids and assign the data points to each centroid to form clusters *C*(*i*), while minimizing the average within cluster dissimilarity, as follows:

1. Randomly initialize *k* cluster centroids *μ*(1),…, *μ*(*k*)
2. For a given cluster assignment *C*, iterate the following steps until the cluster assignments do not change:

a. 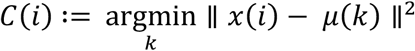.
b. 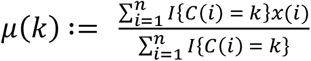.

This specifications make K-means’ performance dependent on initialization of a few hyperparameter, including the number of clusters and the initial cluster centroids.[27,28] Specially, the dependency of K-means on random initialization of the cluster centers often results in the algorithm’s performance being unreliable, when the number of iterations are small.[31] in contrast to K-means, hierarchical clustering is a bottom-up or agglomerative approach that does not require to know the number of cluster in advance. As a result, its performance is not dependent on random initialization of *k* cluster centroids. However, compared with the K-means algorithm, hierarchical clustering algorithm is more computationally intensive.

Hybrid hierarchical-k-means (HK-means) clustering [31] combines the strengths of hierarchical clustering in initializing the cluster centroids, and improves efficiency of the K-means algorithm. The HK-means algorithm relaxes the dependency of K-means algorithm on random initialization of cluster centroids by first computing hierarchical clustering, cutting the tree in *k* clusters, computing the centroids for each cluster, and then using the centroids as the initial cluster centers to run K-means.[31] This hybrid approach accelerates the K-means procedure, and thereby, improves the overall learning.[31]

Performance of the HK-means algorithm still depends on approximation of the number of clusters, *k*. Initializing the number of clusters for the algorithm is a challenging problem, for which a number of ad-hoc (or intuition-based) solutions are available.[32–34] Most of the available solutions do not scale up to large datasets. We used *kluster*, a procedure that uses iterative sampling to produce scalable and efficient cluster number approximation solutions in large datasets.[35]

### 2.4. Implementation

Due to the size of data for each lab, we parallelized the implementation of our clustering solution for identifying the implausible lab observations through the following steps (Figure 2):

1. Extract data on observation *x*, *db*(*x*), from RPDR.
2. Shuffle *db*(*x*) randomly to avoid any specific sorting for parallelization.

To accommodate large datasets, we implemented the clustering approach through parallel commuting.

3. Break *db*(*x*) into *j* folds such that each fold has ***n*** (or fewer) data points.

We controlled the threshold for flagging clusters as implausible 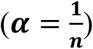 through the selection of the number of data points for parallelization. We experimented with eight thresholds: 1/500, 1/1,000, 1/2,000, 1/3,000, 1/4,000, 1/5,000, 1/6,000, and 1/10,000.

4. Begin parallel computing:

- Extract the subset of *db*(*x*) for fold *j, db*(*x*|*j*).
- Scale *db*(*x*|*j*) and transform the values to the 3^rd^ power – to focus on distribution tails.
- Apply *kluster* procedure[35] to *db*(*x*|*j*) to identify *k* clusters.
- Compute HK-means clustering and assign data points to clusters cluster *C*(1), …, *C*(*k*).
- Count number of data points in each cluster *C*(*k*), *p_k_*.
- Flag all data points in *C*(*k*) as implausible, where *p_k_* ≤ *a*.
5. Produce a report containing all flagged rows.

**Figure 2.**
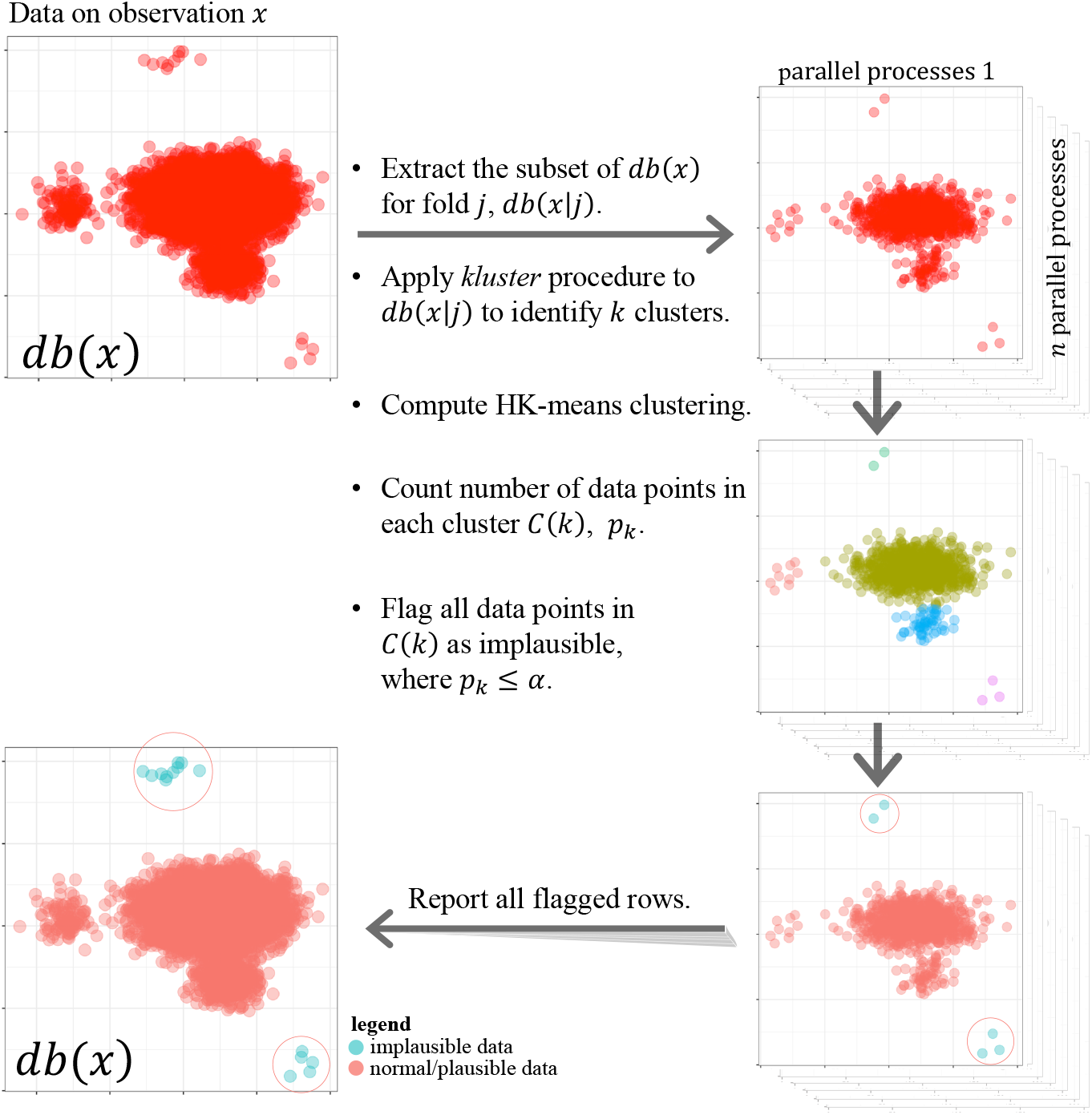
Parallel implementation of the clustering solution for identifying implausible EHR observations.

To measure performance, in this research we compute confusion matrix indices, including false/true positives, false/true negatives, sensitivity, specificity, and fallout.

The R script for implementation of this pseudo code against an i2b2 data model, calculate sensitivity and specificity, and create plots are provided as supplementary materials and also on GitHub (<u>GitHub link to be provided after the peer review</u>).

## 3. Results

After classifying data points into implausible or plausible across the eight αs with our proposed clustering approach, we computed confusion matrix indices against the silver-standard plausibility labels – generated using a combination of expert knowledge, insight from data distributions, and literature search. True positive in the confusion matrix represents the number of truly implausible observations (as identified from the silver-standard labels) that were also identified by the clustering approach as implausible. We calculated both sensitivity and specificity metrics for all implementations (Table 1 in Appendix). Because we were dealing with large datasets, our focus was to reduce false negatives while maintaining high true positive ratios. Therefore, we first evaluated the algorithms based on their specificity (i.e., how many data points identified as plausible by the clustering approach are truly plausible).

### 3.1. Specificity

In all of the 50 lab observations, the clustering approach performed with a septicity greater than 0.9997 (Table 1.a). The best specificities were often obtained from the most stringent ***α*** (1/10,000), which identifies a cluster as implausible only if its population is 1/10,000 of the data subset. In this configuration, we used subsets of 10,000 data points for parallel computing. The 10,000 data points were partitioned into ***n*** clusters, and the cluster with 1 data point was identified as implausible. The lowest specificity, 0.9938, was from the most liberal configuration with ***α*** – 1/500. Overall, we found that specificity increases as ***α*** decreases (Figure 3).

**Figure 3.**
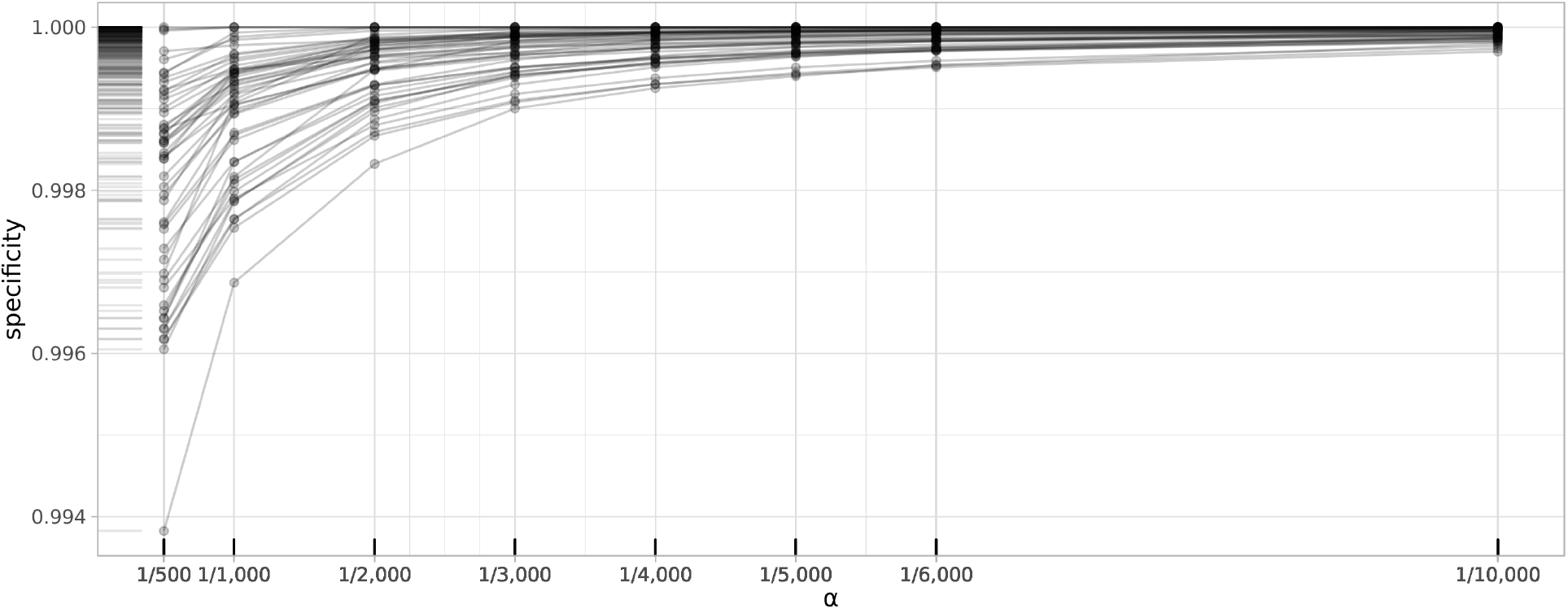
Changes in specificity of the clustering approach by ***α***.

### 3.2. Sensitivity

Sensitivity focuses on true positives, and in our case, represents how many of the implausible observations were picked up by the algorithm. Our sensitivity results we less consistent than the specificities we obtained from the clustering approach. We did not have any implausible observations in 9 of the 50 lab tests – i.e., no true positives in 18 percent of the labs. In the 41 remaining labs, we obtained the best performance from the most liberal configuration ***α***, where 1 in 500 data points was flagged as implausible (Table 1.b). It is important to evaluate the sensitivity results considering the sparsity of positives (implausible observations) in data. The number of implausible observations in the 41 labs ranged from 1 to over 39,000, representing an average of 0.0576 percent of the labs. Considering sparsity, a sensitivity over 0.85 would pick up most of the implausible observations. We obtained sensitivity of over 0.85 in 39 of the 41 labs that had at least 1 implausible observation. For labs, Troponin I.cardiac (LOINC: 10839-9) and Cholesterol in LDL (LOINC: 13457-7), the best sensitivity was 0.0545 and 0.4867, respectively. Troponin I.cardiac (LOINC: 10839-9) was an unusual case with over 39,000 implausible observations, based on our silver standard implausible cutoff of [0-20] and normal range of [0.04-0.39]. Nevertheless, all of the clustering algorithms produced results with 100 percent specificity, meaning that in controversial cases such as Troponin I.cardiac, the clustering approach did not identify any false positive cases. For Cholesterol in LDL (LOINC: 13457-7) there were over 540 positives, or implausible observations, from which the best clustering algorithm detected over 250 as implausible observations.

### 3.3. Comparing the clustering approach with conventional anomaly detection

There are various conventional methods for anomaly detection that are also computationally inexpensive. We evaluated whether our proposed clustering approach performed better than conventional anomaly detection (CAD) methods, namely identifying implausible values using standard deviation and anomaly detection with Mahalanobis distance (Table 2 in Appendix). Overall, the clustering approach produced overwhelmingly better specificity than conventional anomaly detection (Figure 4). Best specificity from CAD methods was obtained from using 6 standard deviations as threshold for identifying outliers, which was outperformed by the clustering approach.

**Figure 4.**
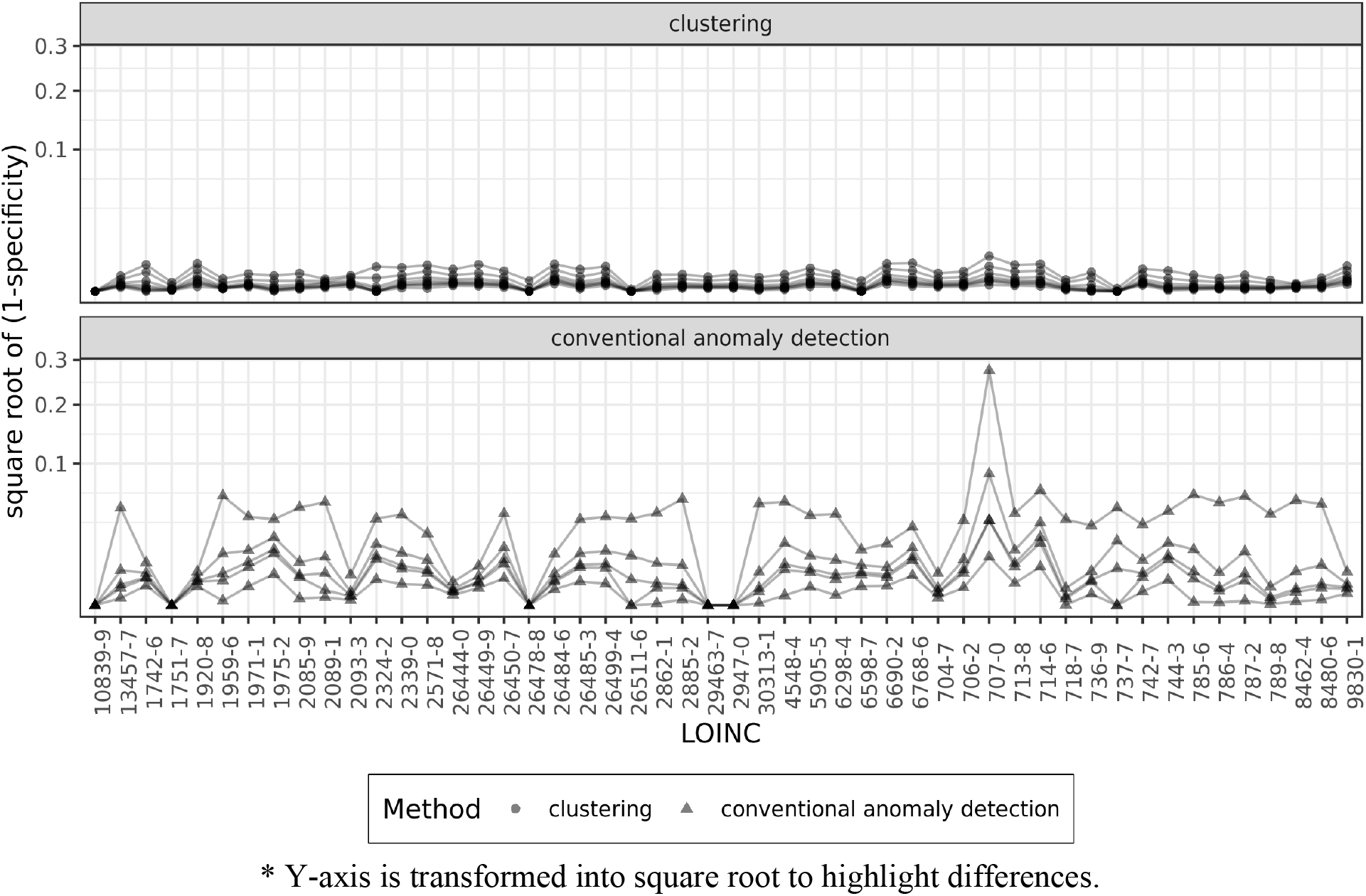
Comparing the specificity (1-specificity) performance between conventional anomaly detection and the clustering approach.

In 31 of the 41 labs, the clustering and CAD produced similar sensitivity (Figure 5). Among CAD methods, the best sensitivity was obtained from applying Mahalanobis Distances and 3.717526 (sqrt of 13.82) as critical value. The conventional anomaly detection (CAD) produced better sensitivity in 9 of the 41 labs for which we had implausible observations. The two largest delta in sensitivity was for Troponin T.cardiac (LOINC: 6598-7), where the clustering approach outperformed the best CAD result by 0.8794, and for Cholesterol in LDL (LOINC: 13457-7), where the best CAD result improved sensitivity by 0.5133. Outside these two labs, the average improvement in sensitivity in 9 labs was 0.1228, which considering the sparsity of implausible observations is minor.

**Figure 5.**
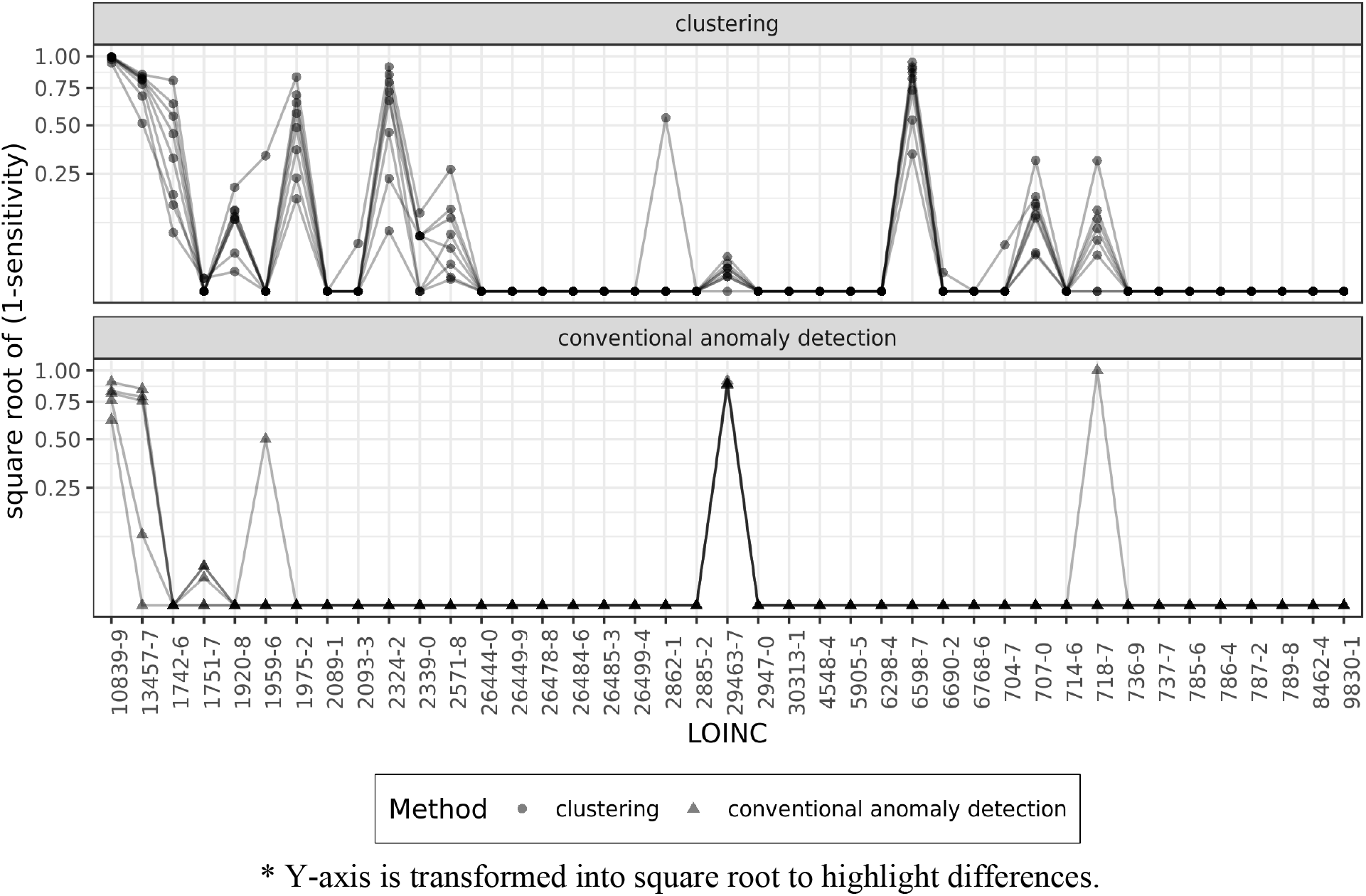
Comparing the sensitivity (1-sensitivity) performance between conventional anomaly detection and the clustering approach.

We further evaluated the differences between CAD methods and our proposed clustering approach through number of false positive cases in each of the labs. As discussed earlier, our goal was to minimize the frequency of observations falsely identified as implausible. Figure 6 illustrates a pairwise comparison of number of false positive cases identified by each approach for each of the 50 lab observations. In 45 out of 50 labs (90 percent), the clustering approach produced a statistically significant smaller number of false positive cases.

**Figure 6.**
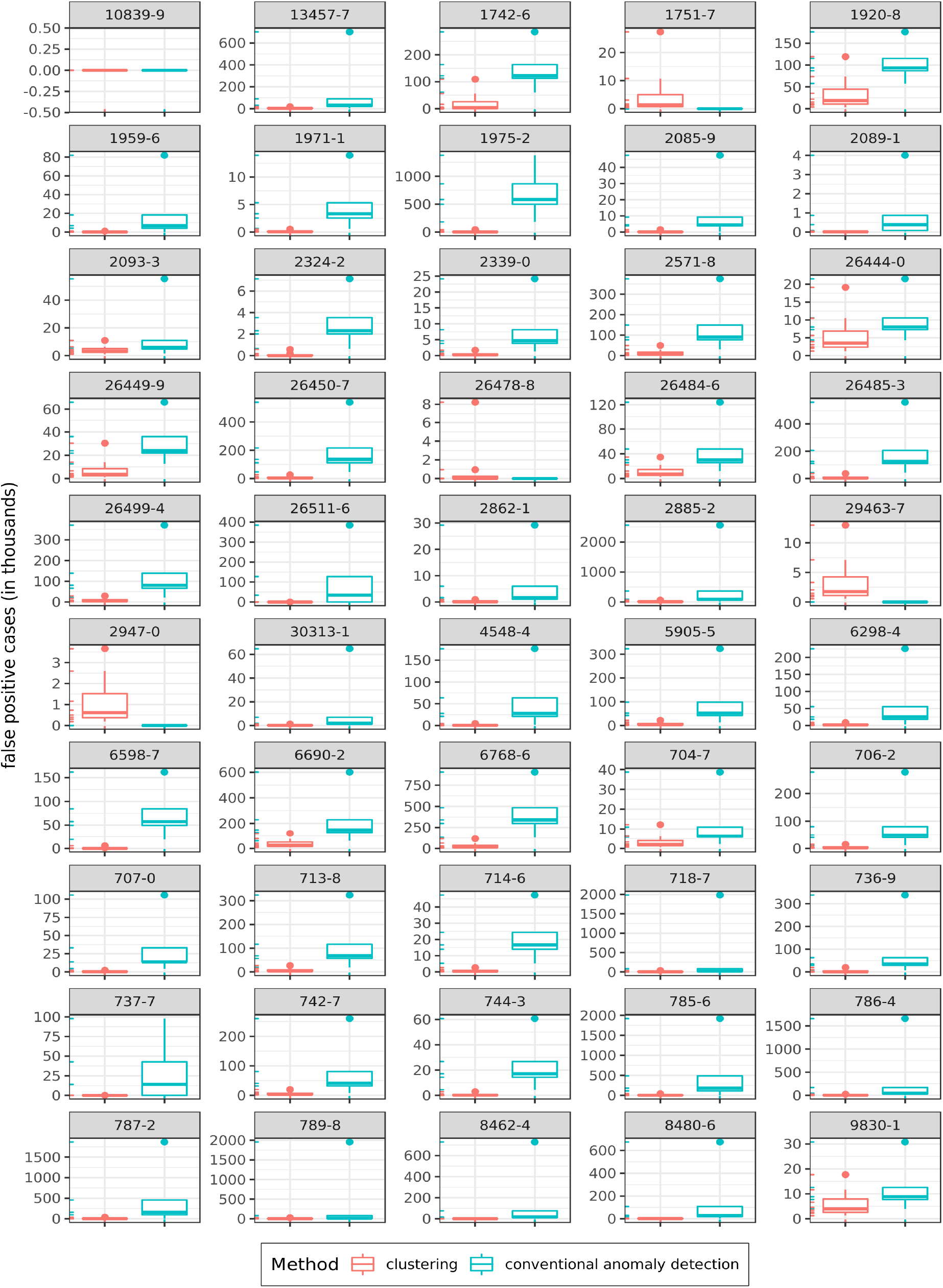
Pairwise comparison of false positive cases between conventional anomaly detection (CAD) and the clustering approaches.

More importantly, when the clustering approach outperformed the CAD approach, the gaps between the two approaches signified a large number of false positives. As the Y-axes show (scale is transformed in thousands) the conventional anomaly detection often identifies thousands of more plausible observations as implausible, compared with the clustering approach.

## 4. Discussion

EHRs provide massive amounts of observational data. Biological data have certain properties that are distinct in their distribution from other types of the so called “Big Data.” We designed, implemented, and tested an unsupervised clustering approach for identifying implausible records in clinical observation data. Our approach is based on two linked hypotheses that 1) if no systematic data entry errors exist, implausible clinical observations in electronic health records are sparse, and therefore 2) if clustered appropriately, clusters with very small populations should represent implausible observations. Using EHR laboratory results data, our results supported both hypotheses. Figure 7 provides an example plot – a set of additional plots on other labs are available in Appendix.

**Figure 7.**
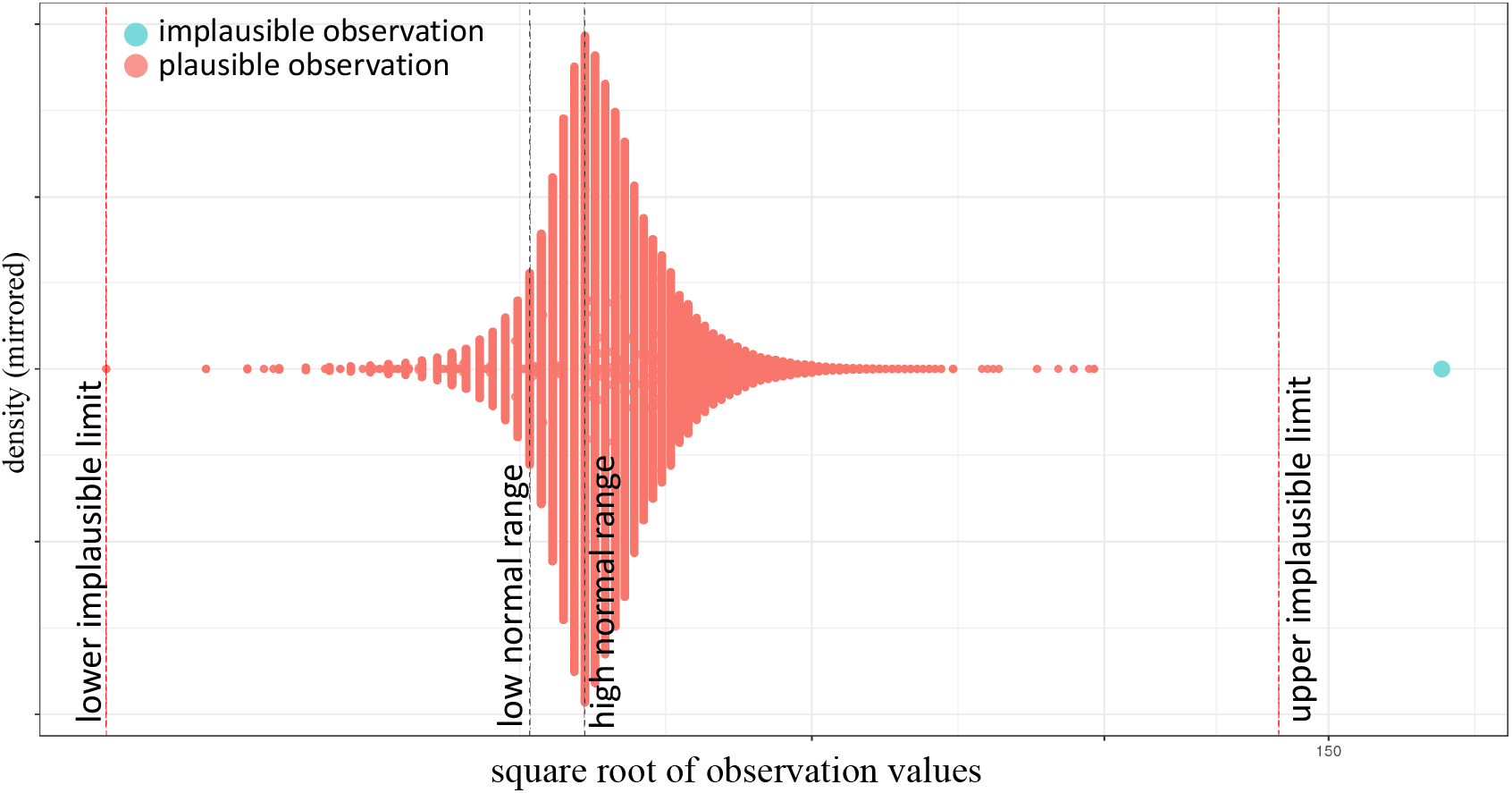
Unsupervised clustering identifies implausible values in Bicarbonate (HC03) (LOINC: 1959-6)

We also demonstrated that the clustering approach outperforms conventional anomaly detection (CAD) approaches in identifying implausible lab observations. In anomaly detection, outliers are often conceived as extreme values at the two tails of the distribution. Our approach expands the conventional definition of an outlier, by searching for observations that sparse considering their value relative to the rest of the data points. Such observations can be found anywhere across the distribution of data – i.e., implausible values can still be extremely high or low, or just different enough from the rest of data points.

The clustering approach for identifying implausible observations offers a precise “Big Data” solution for clinical and biological observations stored in electronic medical records. The clustering process is observation-specific, as the number of clusters and partitioning is specified for each group of observations. As a result, it produced low false positives – plausible observations mistakenly identified as implausible. In contrast, we showed that CAD approaches produce a high number of false positives. This difference is a huge benefit for the clustering approach from an informatics standpoint regarding implementation in large scale data repositories.

### 4.1. Limitations and Directions for Future Research

We used a hybrid hierarchical K-means algorithm, HK-means, because K-means algorithms are generally sensitive to outliers and the hybrid method improves K-means’ reliance on the random centroid initialization. We can imagine that other distance-based or density-based unsupervised clustering algorithms might be also effective in identifying rare implausible clinical observations. Many of clinical observations stored in EHR data are unlabeled. Unsupervised learning approaches offer many promising solutions for patient characterization. Nevertheless, these approaches require specification of several hyper-parameter. In our case, performance of the clustering approach depended on the number of clusters, initial random assignment of cluster centroids, and the threshold for flagging clusters as implausible. In addition, implementation of the approach on very large datasets was challenging. To address some of these challenges, we had to be creative in selecting the clustering algorithm, applying feature transformations, and developing the *kluster* procedure to approximate the number of clusters, all of which would be practical for future unsupervised learning efforts that aim to ascertain meaningful patient sub-groups.

We randomly shuffled the data to prevent any potential sorting in breaking the data to folds for parallel computing. Further research is needed to ascertain whether some systematic approach to parallelization (e.g., breaking the data by age and gender) can improve the unsupervised implausible observation detection results.

Moreover, we have tested the clustering approach against 50 EHR observations. This demonstration was limited to a small set of observations due to the need for silver-standards to measure sensitivity and specificity of our proposed approach. However, we encourage the readers to envision further applications of this approach to other clinical observations, as well as complex combinations of observations in multiple dimensions, for which specification of manual silver-standards are virtually implausible.

### 4.2. Implementation Considerations

The algorithmic solution we presented in this paper proved as a feasible alternative for replacing the current manual rule-based procedures for identifying biologically implausible values. Given the size of data and the emphasis on sensitivity versus specificity, the choice of ***α*** (the ratio for flagging a cluster as implausible) can vary. In large clinical data repositories, an ***α*** between 1/4,000 and 1/,6000 would provide good balance between true positives and false negatives. A smaller ***α*** is also computationally more expensive. When the size of the dataset is small, a small ***α*** will be more appropriate. Due to superb specificity implementing the clustering approach offers a low-risk solution to an expensive manual procedure that is hard to implement. We envision the proposed solution to constantly operate on the data base servers where EHR data are stored. We recommend that our algorithmic solution should be used to augment (rather than replace) human decision-making for improving quality of EHR data. After the implausible values are detected and flagged, a workflow is needed to initiate further actions needed in order to determine the destiny of the flagged observations. As we showed, because the flagged records are also sparse, the frequency of such flags should not be of concern.

## 5. Conclusion

Detecting implausible clinical observations in Electronic Health Record (EHR) data is a challenge, requiring availability of standards thresholds and rule-based procedures to query observations that are out of the plausibility range. Establishing rule-based procedures to address this task entails extensive hard-coding that would accumulate over time and dimensionality. We proposed an alternative viable algorithmic solution, using unsupervised clustering approach. The clustering approach is superior than conventional anomaly detection approaches and adaptable to different types of numerical EHR observation data.

